# Machine learning applied to predicting microorganism growth temperatures and enzyme catalytic optima

**DOI:** 10.1101/522342

**Authors:** Gang Li, Kersten S. Rabe, Jens Nielsen, Martin K. M. Engqvist

## Abstract

Enzymes that catalyze chemical reactions at high temperatures are used for industrial biocatalysis, applications in molecular biology, and as highly evolvable starting points for protein engineering. The optimal growth temperature (OGT) of organisms is commonly used to estimate the stability of enzymes encoded in their genomes, but the number of experimentally determined OGT values are limited, particularly for thermophilic organisms. Here, we report on the development of a machine learning model that can accurately predict OGT for bacteria, archaea and microbial eukaryotes directly from their proteome-wide 2-mer amino acid composition. The trained model is made freely available for re-use. In a subsequent step we OGT data in combination with amino acid composition of individual enzymes to develop a second machine learning model – for prediction of enzyme catalytic temperature optima (*T*_*opt*_). The resulting model generates enzyme *T*_*opt*_ estimates that are far superior to using OGT alone. Finally, we predict *T*_*opt*_ for 6.5 million enzymes, covering 4,447 enzyme classes, and make the resulting dataset available for researchers. This work enables simple and rapid identification of enzymes that are potentially functional at extreme temperatures.

## 1 Introduction

Enzymes that remain active at high temperatures, sometimes referred to as thermozymes, are used to catalyze chemical reactions in industrial processes (*1–8*), for applications in molecular biology (*9–14*), and for providing highly evolvable starting points for protein engineering (*15–19*). When testing new enzymes for these applications the optimal growth temperature (OGT) of microorganisms is commonly used to estimate protein stability – enzymes derived from thermophilic organisms are expected to be both stable and active at high temperatures.

Although frequently successful, using OGT as an estimate faces two challenges. First, for many microorganisms with experimentally determined OGT this information is not readily accessible. This challenge has been partially addressed through the creation of public databases and datasets (*20–23*). However, the OGT for the vast majority of microbial organisms is currently unknown since determining the OGT of a microorganism is a laborious process that requires cultivation in temperature-controlled conditions. The number of microorganisms that can be cultured in the laboratory is only a small fraction of the total diversity in nature(*24*). Consequently, many suitable enzyme catalysts likely remain untested and undiscovered. Second, using OGT to estimate enzyme catalytic optima (*T*_*opt*_) constitutes a rough approximation, with many enzymes displaying *T*_*opt*_ at temperatures significantly higher or lower than the OGT (*22,25*). The Pearson correlation between the *T*_*opt*_ of individual enzymes and OGT is only 0.48(*22*). In practice this means that enzymes from thermophilic organisms may be optimally active at significantly lower temperatures than expected.

Due to these challenges a simple way to computationally estimate (1) the OGT of microbes and (2) the *T*_*opt*_ of enzymes is in demand. For such computational estimations to be feasible there must be general trends for how quantifiable biological properties change with growth temperature, there must be a signal that can be modeled. The OGT of microorganisms is an important physiological parameter that has been widely used to understand the strategies organisms use to adapt their genomes and proteomes to different environmental conditions(*26–28*). Many genomic and proteomic features that are strongly correlated with OGT have been revealed. Examples include the existence of thermophile-specific enzymes(*29*), the presence or absence of certain dinucleotides(*30*), the GC content of structural RNAs(*31*), as well as amino acid composition of the proteome(*26,32*). Examples such as these indicate that estimating OGT directly from genomes or proteomes may indeed be feasible.

Statistical tools, such as regression and classification, have been used to model the correlation between OGT and biological features. For example, the OGT of 22 bacteria could be predicted using a linear combination of either dinucleotide or amino acid composition(*30*). Additionally, Zeldovich found that the sum fraction of the seven amino acids I, V, Y, W, R, E and L showed a correlation coefficient as high as 0.93 with OGT in a dataset consisting of 204 proteomes of archaea and bacteria(*32*). Jensen et al developed a Bayesian classifier to distinguish three thermophilicity classes (thermophiles, mesophiles and psychrophiles) based on 77 bacteria with known OGT(*33*). Training datasets containing the OGTs for a large number of organisms have been hard to obtain, something which has prevented the development of state-of-the-art machine learning models for OGT prediction.

While there are only a few published models predicting organism OGT, we know of no computational tools to estimate *T*_*opt*_. However, many methods for the estimation of protein stability have been developed. We wish to emphasize the difference between these two measures as stability is an indication of the folding state of the protein, without information regarding catalytic activity, whereas *T*_*opt*_ implicitly assumes stability and instead indicates the temperature of optimal catalysis. Methods for predicting protein stability fall into two main categories; predicting the stability of whole proteins, and predicting the stability change in a protein upon amino acid substitutions. Machine learning has been used extensively for the prediction of stability change upon amino acid substitutions(*34–38*), while only a few methods have been developed for the prediction of stability of whole protein empirically(*39–42*). However, computational prediction of protein stability is challenging since it usually needs an accurate calculation of Gibbs-free energy change of protein unfolding process(*41,42*), which relies mainly on high-quality protein structures. Such structures are limited in number, thereby reducing the applicability of these methods for identifying thermostable enzymes for industrial applications.

Here, we address the challenge of identifying proteins active at high temperatures in in three steps. First, we build a machine learning model to accurately predict OGT using features extracted from all proteins encoded by an organism’s genome. This model is used to assign OGT values for organisms without experimental data. Second, we significatly improve the prediction of enzyme *T*_*opt*_ values by using OGT in combination with sequence information of individual enzymes. Those predictions are significantly more accurate than using OGT alone for the prediction. Finally, we make use of the predictive models to estimate *T*_*opt*_ for 6.5 million enzymes, covering 4,447 enzyme classes in the BRENDA(*43*) database. The OGT model and enzyme *T*_*opt*_ estimates are made freely available for reuse (https://github.com/EngqvistLab/Tome).

## 2 Methods

### 2.1 Software

All machine learning analysis were conducted with scikit-learn package (version 0.19.1)(*44*) using Python version 2.7.14. The module and model hyperparameters used are listed in Supplementary Table S2. Python code for proteome analysis, machine learning and data visualization are available from the authors upon request. The source code for the Tome package is available under a permissive GPLv3 license at GitHub (https://github.com/EngqvistLab/Tome).

### 2.2 Proteome dataset

The bulk of protein sequence data used in this work was obtained from Ensembl Genomes release 37, obtained in September 2017 (http://ensemblgenomes.org/). For all archaea and bacteria listed at ftp://ftp.ensemblgenomes.org/pub/bacteria/release-37/fasta/ fasta files containing protein sequences were downloaded. Similarly, fasta files containing protein sequences for all fungi listed at ftp://ftp.ensemblgenomes.org/pub/fungi/release-37/fasta/ were downloaded. As a complement to the Ensembl Genome data we made use of protein data from RefSeq release 87, obtained in March 2017 (https://www.ncbi.nlm.nih.gov/refseq/). Fasta files containing a nonredundant set of protein sequences for each organism were downloaded from ftp://ftp.ncbi.nlm.nih.gov/refseq/release/ for archaea, bacteria, fungi and protozoa.

In many cases the Ensembl Genomes and RefSeq datasets both contained information for the same organism, or for several strains of the same organism. Therefore, to combine the two datasets, the following steps were followed: First, where multiple strains from the same organism were present in the Ensembl Genomes dataset, the strain with the largest file size, indicating the greatest number of amino acids in the downloaded fasta file, was selected for analysis. Other strains for that organism were discarded. Second, where the same organism was present in both the Ensembl Genomes and RefSeq datasets the one from Ensembl Genomes was retained and the one from RefSeq was discarded. In this way a protein dataset comprising protein sequence data for 7,565 microorganisms was obtained. Of these 5,325 originated from Ensembl Genomes and 2,240 originated from RefSeq.

For each organism in the protein dataset we attempted to annotate it with its optimal growth temperature. In this annotation procedure organism names were stemmed to the species level (ignoring strain designations) and cross-referenced with a published dataset containing growth temperatures for 21,498 microorganisms (https://doi.org/10.5281/zenodo.1175608). Growth temperatures could be associated with the protein sequence data from 5,762 organisms, whereas 1,803 were left unannotated.

### 2.3 Estimation of threshold

For each proteome, the total length of each protein was calculated. Then the amino acid frequencies and the total number of residues of the first n proteins (n = 1, 2, ..N, were N is the total number of proteins) were calculated sequentially. The data points in the last one-third of all residues added were used to measure the stability of the calculated amino acid frequencies. Three different metrics were designed: (1) the absolute slope value *|a*_i_*|* in the linear regression between the number of residues and amino acid frequency; (2) frequency variance of these selected frequencies 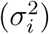 and (3) varying range (*R*), the difference between maximal frequency and minimal frequency. Ideally, 0 was expected for all these three metrics if there is an absolutely stable amino acid frequency in a given proteome. Finally, for each proteome, the maximal 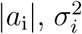 and *R* of 20 amino acids of each proteome (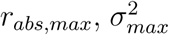 and *R*_*max*_) were used to measure whether frequencies were stable.

To test the effect of the protein order in a proteome in the above analysis, a shullfing strategy was applied. Firstly, equal coverage over the log_10_-transformed proteome size range 3-7.5 was ensured by performing the random sampling in 20 bins. One proteome was randomly selected for each bin and this resulted in 17 selected proteomes as there is no proteome in 3 of these bins. The order of the proteins in each proteome was randomly shuffled and then 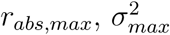 and *R*_*max*_ were calculated. Each proteome was shuffled for 100 times.

### 2.4 Machine learning workflow for OGT model

20 amino acid frequencies and 400 dipeptide frequencies were extracted for each proteome. Then, each of these features were normalized by 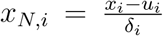, where *x*_*i*_ is the values of feature *i, u*_*i*_ and *δ*_*i*_, are mean and standard derivation of *x*_*i*_, respectively. The following six models were selected and their performance were tested on the annotated and filtered proteome dataset using single amino acid frequencies (AA), dipeptide frequencies (Dipeptide) or the two together (AA+Dipeptide): Linear regression (Linear), bayesian ridge, elastic net, decision tree, support vector regression (SVR) and random forest. 5-fold cross-validation was used for the calculation of *R*^2^ scores. For SVR, elastic net, decision tree and random forest models, an additional 3-fold internal cross-validation were used to optimize the hyperparameters. The model with the highest *R*^2^ score was selected and trained, without cross-validation, on the whole dataset. For the prediction of OGT for those un-annotated organisms, dipeptide frequencies were normalized by 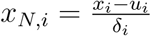, where *x*_*i*_ is the values of feature i. *u*_*i*_ and *δ*_*i*_, are mean and standard derivation of feature *i* in the training dataset, respectively.

### 2.5 OGT Model validation

For validating the OGT prediction model we sampled 54 species with predicted growth temperatures (for which no growth temperatures were available in the original dataset) at random. Equal coverage over the temperature range 0-100°C was ensured by performing the random sampling in 10 bins, each spanning a 10°C temperature range. The primary scientific literature was then manually searched to obtain documented experimental growth temperatures for the sampled organisms. For 45 organisms a documented growth temperature could be found, for 9 organisms it could not. The accuracy of predicted OGT was assessed by computing the Pearson correlation with experimental OGT.

In a second approach to validating the OGT prediction model we used Python scripts and the Zolera SOAP package (https://pypi.python.org/pypi/ZSI/) to extract all available experimentally determined enzyme temperature optima from the BRENDA enzyme database https://www.brenda-enzymes.org/ release 2018.2 (July 2018). Data coming from the same enzyme was de-duplicated by averaging temperature optima from records with the same EC number and originating from the same organism. For each organism with catalytic optima for more than five enzymes the arithmetic mean of those optima were calculated. Those organisms present in both the BRENDA enzyme data as well as the dataset with predicted OGT were identified through cross-referencing species names. The accuracy of predicted OGT was assessed by computing the Pearson correlation between predicted OGT and mean catalytic optima of enzymes.

### 2.6 Machine learning workflow for *T*_*opt*_ model

UniProt identifiers for proteins with an experimentally determined catalytic optimum were obtained from the “TEMPERATURE OPTIMUM” table in the web pages of the BRENDA database, release 2018.2 (July 2018). These identifiers were filtered to retain only those associated with an organism with experimentally determined OGT. After further filtering to remove sequences containing “X” (unknown amino acid), a dataset with 2,609 enzymes was generated. The protein sequences for each of these identifiers were downloaded from the UniProt database in fasta format.

The following features were extracted for each enzyme: (1) 20 amino acid frequencies (AA); (2) 400 dipeptide frequencies (Dipeptide); (3) OGT of its source organism; (4) Basic features including protein length, isoelectric point, molecular weight, aromaticity(*45*), instability index(*46*), gravy(*47*) and fraction of three secondary structure units: helix, turn and sheet. These features were extracted with the module Bio.SeqUtils.ProtParam.ProteinAnalysis in Biopython(*48*) (version 1.70). Additionally, siix binary features were extracted: EC=1, 2, 3, 4, 5, 6. These numbers represent the first digit in a EC number. All features except binary features were normalized as described in section “Machine learning workflow for OGT model”. The following five models were tested on the resulting dataset: bayesian ridge, elastic net, decision tree, support vector regression (SVR) and random forest. The linear model was not used due to its poor performance on any datasets containing dipeptide frequences (negative *R*^2^ scores by cross-validation). The performance of the five regression models was tested using the same cross-validation strategy as for OGT. In addition, to test the accuracy of using OGT of the organism as an estimation of enzyme *T*_*opt*_, the *R*^2^ score between each enzymes *T*_*opt*_ and associated OGT was calculated. The model with the highest *R*^2^ score was chosen and trained on the full training dataset.

### 2.7 BRENDA annotation

Protein sequence data for each EC class was obtained by downloading comma-separated flatfiles from the BRENDA database version 2018.2 (July 2018). Each sequence in these files contain information regarding source organism as well as unique UniProt identifiers. Where possible, each protein sequence was associated with an OGT value by mapping the source organism name to the OGT dataset from https://doi.org/10.5281/zenodo.1175608. Those sequences were firstly mapped to the existing *T*_*opt*_ values in BRENDA by matching EC-UniProt id pair. For those enzymes without any experimental *T*_*opt*_ values, the amino acid frequencies were calculated (ignore all “X” in the sequence). All 20 amino acid frequencies as well as the OGT variable were normalized by 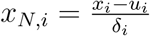, where *x*_*i*_ is the values of feature i. *u*_*i*_ and *δ*_*i*_, are mean and standard derivation of feature *i* in the original training dataset, respectively. Finally, the normalized values were used for the prediction of *T*_*opt*_ by the previously generated random forest regressor trained on the AA+OGT datasets. The predicted enzyme *T*_*opt*_ and annotated OGT values of these enzymes are freely available for download and re-use (https://zenodo.org/record/2539114, https://doi.org/10.5281/zenodo.2539114).

## 3 Results and discussion

### 3.1 Collection of optimal growth temperature and proteomes of microorganisms

Protein amino acid composition is strongly correlated with OGT(*30,32*). For this reason we decided to train machine learning models using the amino acid composition as features. To build such a model we first established a training dataset. To this end, we downloaded an OGT dataset (https://doi.org/10.5281/zenodo.1175608), which contains data for 21,498 microorganisms, including bacteria, archaea and eukarya(*22*). Using this dataset, all proteins from 5,761 organisms from RefSeq (https://www.ncbi.nlm.nih.gov/refseq/) and Ensembl genomes (http://ensemblgenomes.org/) could be associated with an OGT value (we refer this as the annotated dataset), while proteins from an additional 1,803 organisms could not be associated with an OGT value (we refer this as the unannotated dataset) (Figure 1).

**Figure 1:**
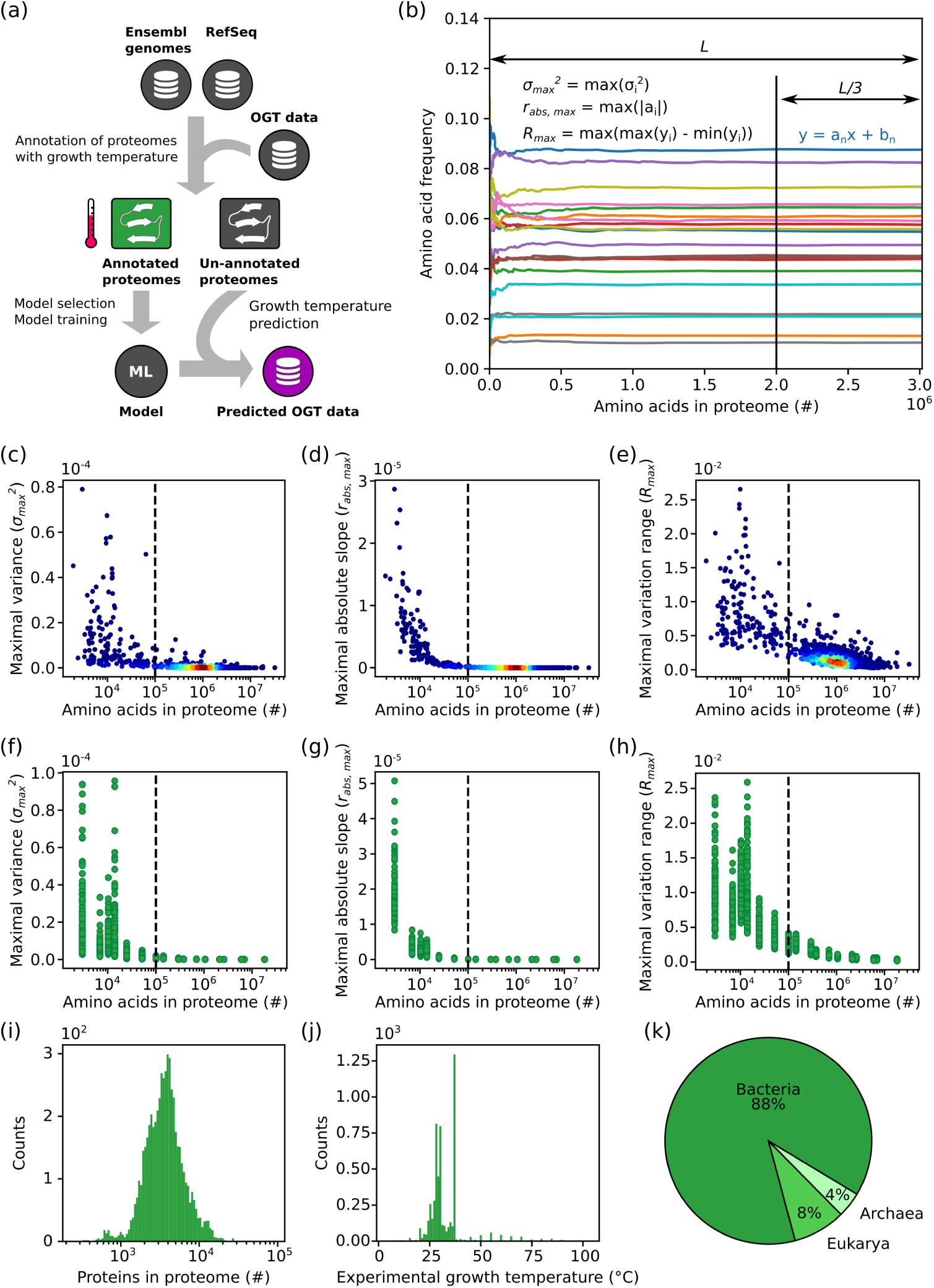
Variability in amino acid frequencies decrease log-linearly with proteome size. (a) Schematic overview of process to build a machine learning model to predict OGT. Protein records from Ensembl genomes (bacteria and fungi) and RefSeq (bacteria, archaea and fungi) were downloaded. Sequences were annotated with the growth temperature of the organism from which they originate. Sequences from organisms that could not be annotated, i.e. for which there is no available information about the OGT for the organism, were retained in a separate un-annotated dataset. Amino acid frequencies of the annotated sequences were used to train a statistical model. This model was in turn used to predict growth temperatures for the un-annotated dataset. OGT: optimal growth temperature. (b) The frequency of each amino acid was plotted against the number of amino acids used to calculate the frequency. The final third part was fitted to a linear model to get the absolute slope value (*|a*_i_*|*), as well as its frequency variance 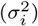 and varying range (*R*). The maximal 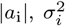 and *R* of 20 amino acids of each proteome (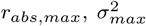 and *R*_*max*_) give measures of whether frequencies were stable. The calculated (c) 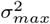, (d) *r*_*abs,max*_ and (e) *R*_*max*_ of all species in the dataset were plotted against the number of amino acids in the proteome. The dashed line indicates the cutoff for the selection of proteomes based on size. Effect of protein order on (f) 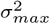, (g) *r*_*abs,max*_ and (h) *R*_*max*_. 17 proteomes with different size were randomly selected. Proteins in each proteome were shuffled 100 times and the three metrics for each shuffled proteome were calculated. (i) Distribution of proteome sizes in the annotated dataset after filtering. (j) Distribution of growth temperatures in the annotated dataset after filtering. (k) Proportion of species belonging to the three different taxonomic superkingdoms in the filtered dataset.

For each organism in both the annotated and unannotated dataset we calculated the global amino acid monomer and dipeptide frequencies. However, some organisms in the dataset contain only a small number of protein sequences, as a consequence the amino acid composition obtained from those sequences may not represent the true amino acid composition of the complete proteome. To address this problem we applied a filtering step. As it was unclear how many protein sequences are required to obtain a stable amino acid composition we designed three different metrics (see Methods for details) to test how much protein sequence data was needed to obtain a stable amino acid composition. (Figure 1b). For each organism in the annotated dataset the three metrics were calculated for every protein sequence added in order to observe at which point the values stop fluctuating. Using this analysis on amino acid monomer frequencies we found that at least 10^5^ amino acids are needed to get a stable amino acid composition (Figure 1c, d, e). Repeating this analysis for amino acid dipeptides resulted in the same threshold (Figure S1).

A further concern was that the order in which proteins appear in the input files may affect our cutoff analysis. For this reason, proteins from 17 organisms with different sizes of available proteomes were randomly selected. For each of these organisms the order in which protein sequences appear was shuffled and the three metrics were calculated. The shuffling, with subsequent analysis, was repeated 100 times. As expected, the analysis shows a high initial variability, where few sequences have been analyzed, but with increasing numbers of averaged proteins the values stabilized and converged (Figure 1f, g, h). From this analysis it is clear that the arrangement of proteins in a proteome has a negligible effect when the proteome size is larger than 10^5^, and we therefore only chose organisms with at least 10^5^ amino acids in the dataset for further analysis. This approach resulted in a training dataset with 5,532 organisms annotated with OGT, as well as a dataset with 1,438 un-annotated organisms. The annotated training dataset comprises 4,974 bacteria, 222 archaea and 337 eukarya (Figure 1) and is much larger than those used in other approaches, such as 22 bacteria(*30*), 77 bacteria(*33*) or 204 prokaryotes(*32*). In the annotated dataset the number of proteins in each organism follows a normal distribution centered around 3,000 (Figure 1i). The OGT distribution is, however, highly skewed with the majority of organisms having an OGT in the range 25-30°C and at 37°C (Figure 1j). The number of organisms in the data set with an OGT higher than 40°C is 425 (340 for higher than 50°C.

### 3.2 OGT can be accurately predicted from amino acid composition of the proteome

For each organism in the annotated dataset we calculated the global amino acid monomer frequencies (20 features) as well as amino acid dipeptide frequencies (400 features). To get the best feature set and statistical model for the prediction of OGT, we tested six different regression models and compared their performance on the monomer dataset and the dipeptide dataset. As shown in Figure 2a, a 5-fold cross-validation was applied to evaluate the performance of different regression models. Using the 20 amino acid frequencies, non-linear models (SVR and Random forest) perform much better than linear models (Linear, Elastic net, Bayesian ridge regression). The superior performance of non-linear models suggests that there are important non-linear relationships between amino acid frequencies and OGT. In contrast, all models except decision tree show an almost identical performance when using dipeptide frequencies. Since models trained on each of the two datasets individually show good performance we reasoned that models trained on the combined datasets may be even better performing. However, contrary to this expectation the six models trained using both monomer dataset and dipeptide dataset together do not show improvement (Figure 2a).

**Figure 2:**
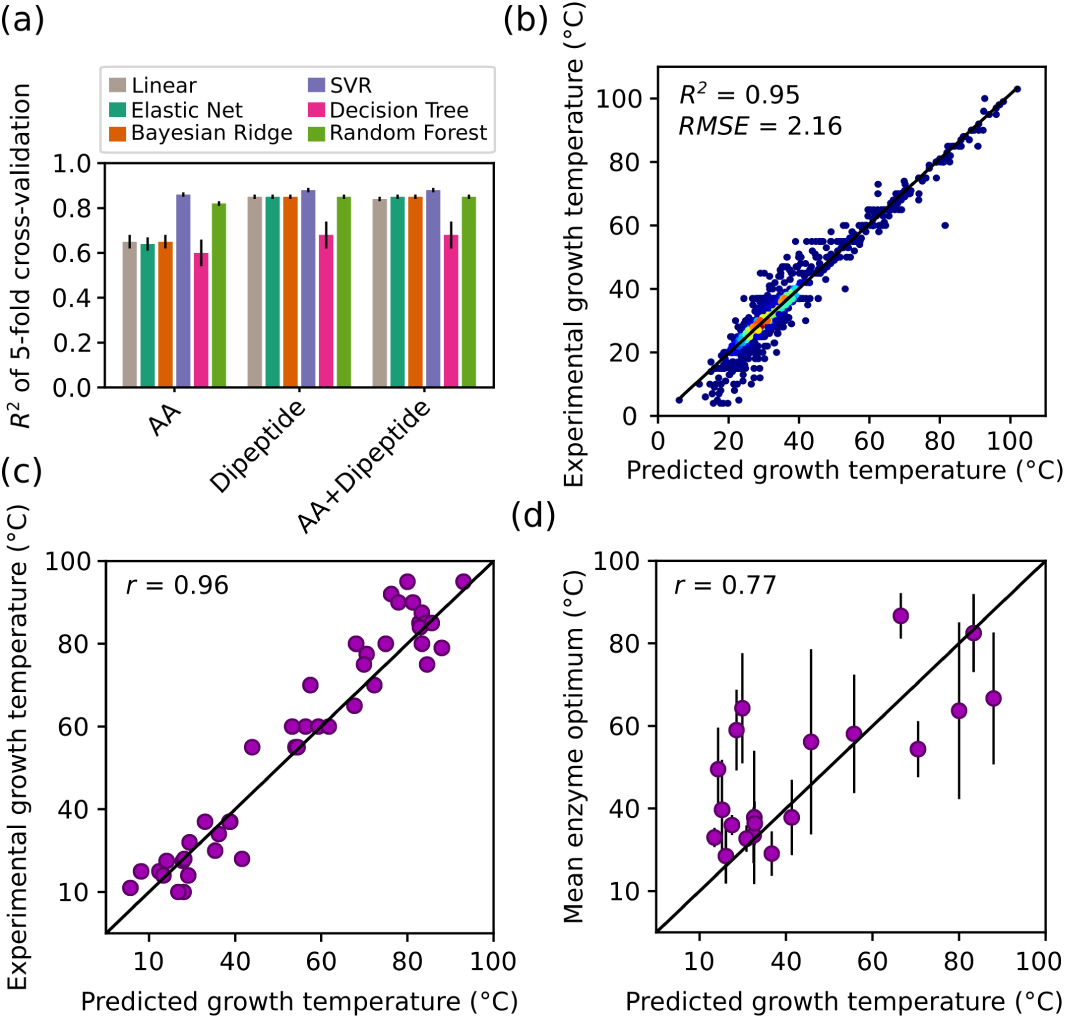
Model development for OGT prediction. (a) *R*^2^ score obtained by a 5-fold cross-validation for six different regression models. Error bars represent the standard derivation of *R*^2^ scores. (b) Performance of the final SVR (support vector regression) model trained on dipeptide data. The correlation between predicted organism growth temperatures and those present in the original annotated dataset was evaluated. RMSE: root mean square error. Colors indicate the density of the points. (c) Correlation between literature values for growth temperatures and predicted growth temperatures. Species for unannotated dataset were sampled at random, but with ensuring equal coverage over the temperature range. Growth temperatures for these organisms were obtained by manually searching the primary scientific literature. (d) Correlation between the mean enzyme temperature optima and predicted growth temperatures for each species present in both datasets. Only organisms with optima for at least five enzymes are shown. Error bars show the standard deviation. In (c and d) *r* denotes Pearson’s correlation coefficient.

A final SVR model was trained on the whole dipeptide dataset, without cross-validation, and stored for further use. This model can explain an astounding 95% (88% by cross-validation) of the variance in OGT (Figure 2b). This model has significantly higher predictive accuracy than other published models (Figure S2 and Figure S3. We propose that the high predictive accuracy results from two features of our approach; the size and quality of the training data used, and the use of non-linear regression models. As a direct consequence of the increased size of the dataset, we could train models that are more general applicable. We find that in general non-linear models outperform linear models when using amino acid frequencies (Figure 2a). This suggests that the linear models used previously such as that from Nakashima et al.(*30*) might be further improved by non-linear regression to correlate the amino acid frequencies to OGT.

### 3.3 Validation of the SVR model for growth temperature prediction

Leveraging the final SVR model, the OGT of 1,438 organisms in the unannotated dataset were predicted (Figure 1a). These OGT predictions were validated using two separate approaches. First, we performed a manual literature search to find experimentally obtained OGTs for a subset of the organisms (for which no experimental OGT was present in our original dataset). We randomly sampled 54 of the organisms with predicted OGTs, in a manner that ensured even spread across temperatures. For 45 of the 54 organisms, OGT values could indeed be found in published peer-reviewed articles (Table S1). The agreement between the predicted OGT and the ones collected from literature is very high, with a Pearson correlation coefficient of 0.96 (Figure 2c). Second, we seized on the fact that the average temperature optimum of catalysis (*T*_*opt*_) of at least five enzymes from an organism shows a Pearson correlation above 0.75 with growth temperature(*22*). Of the 1,438 organisms with predicted OGT only 23 were found to have at least five enzymes with *T*_*opt*_ available in BRENDA. Plotting the arethmatic mean of these enzyme optima against the predicted OGT for each organism reveals a strong correlation, with a Pearson’s correlation coefficient of 0.77 (Figure 2d). Indeed, this correlation is the same as that obtained with experimentally determined organism OGTs(*22*), again showing that the predicted OGTs are very accurate.

### 3.4 Improved estimation of enzyme temperature optima using machine learning

In biotechnology and protein engineering OGT is typically used directly to guide the discovery of thermostable enzymes(*3,21*). We hypothesized that the accuracy of this estimation could be improved by also considering enzyme sequence information in a machine learning framework. A training dataset was generated by collecting 2,609 enzymes that: (1) have *T*_*opt*_ and protein sequence data in the BRENDA database, and (2) come from organisms with an experimentally determined OGT (Figure 3a, b). We first tested the accuracy of directly using OGT as an estimation of *T*_*opt*_ and found that only 25% of the enzyme *T*_*opt*_ variance could be explained (Figure 3c, black bar). Then, to improve the accuracy of this estimation using machine learning, we extracted three feature sets from the enzyme sequences, namely amino acid frequencies, dipeptide frequencies and other basic protein properties like length, isoelectric point etc. (See Methods). Six regression models were trained and tested on these feature sets individually, as well as the two and three sets combined, with a 5-fold crossvalidation approach. As shown in Figure 3c, the best model (SVR) trained on amino acid frequencies achieved a slightly improved accuracy compared to OGT, as quantified by an *R*^2^ score of around 30%. Using dipeptide frequencies alone in combination with amino acid frequencies did not further improve the accuracy.

**Figure 3:**
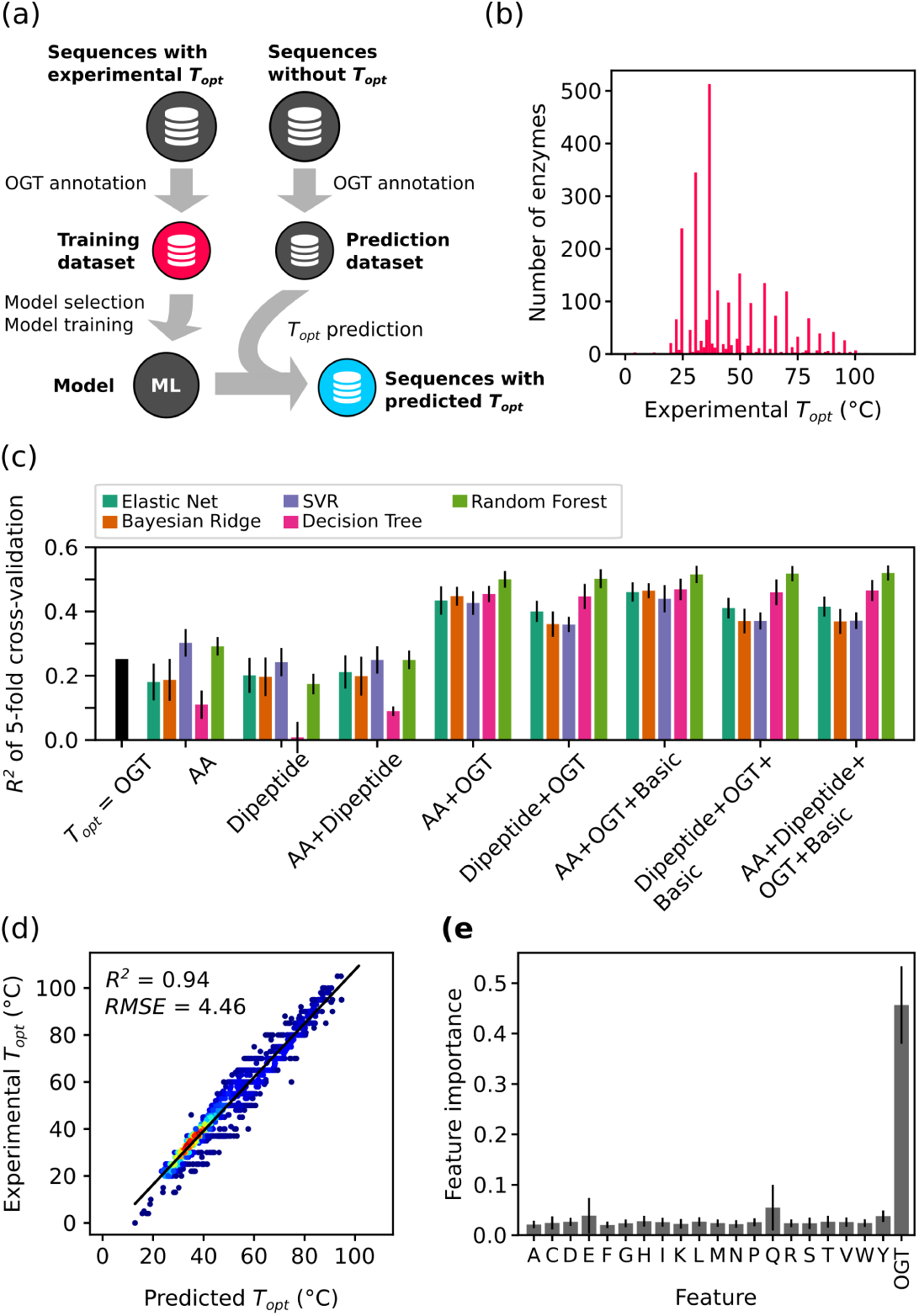
Model development for prediction of enzyme temperature optima. (a) Schematic overview of process to build a *T*_*opt*_ prediction model. (b) The distribution of enzyme temperature optima in training dataset. (c) 5-fold cross-validation results for five regression models on different feature sets. The *T*_*opt*_ = OGT bar shows the explained variance when using OGT as the estimation of enzyme *T*_*opt*_. Error bars shows the standard deviation of *R*^2^ scores obtained in 5-fold cross validation. AA, amino acid frequencies; Dipeptide, dimer frequencies; OGT, optimal growth temperature of source organism; Basic, basic information of proteins, like length, isoelectric point etc., see details in Methods section. (d) Performance of the final random forest model trained on AA+OGT data. The correlation between predicted and experimental *T*_*opt*_ was evaluated. RMSE: root mean square error. Colors indicate the density of the points. (e) The feature importance in the final random forest model. Error bars indicates the standard deviation of feature importances of 1,000 estimators.

Since OGT and sequence-derived features each produce estimates of similar accuracy (25% and 30%, respectively) we tested whether their combined use could boost predictive power. In line with our original hypothesis the best model (random forest) trained on the combination of amino acid frequencies and OGT almost doubled the model predictive accuracy to over 50%. Further inclusion of other basic enzyme properties (see Methods) did not further improve the accuracy (Figure 3c).

To generate a final model for the prediction of enzyme temperature optima the random forest model was re-trained – without cross-validation – using the full set of amino acid frequencies and OGT data (Figure 3d). In this final model, OGT of the source organism is the most informative individual feature, whereas the 20 amino acid frequencies combined contribute over half of the predictive power of the model (Figure 3e). This is a remarkable result that demonstrates a clear importance of combining physiological parameters, such as OGT, with sequence information in the estimation of protein properties. We speculate that using larger training datasets and extracting more descriptive features (both from sequence and physiological parameters) in conjunction with advanced machine learning models, like deep learning(*49*), may further improve the prediction of enzyme *T*_*opt*_. The *R*^2^ score of 51% obtained for *T*_*opt*_ predictions in this study could be used as a benchmark accuracy for future model development.

### 3.5 Annotating enzymes in BRENDA using OGT and predicted *T*_*opt*_

Currently, a main resource for enzyme data is the BRENDA database(*43*). However, there are approximately 12 million native protein sequences in BRENDA while there are only about 33,000 *T*_*opt*_ records, many of which are not connected to a protein sequence. We made use of the *T*_*opt*_ prediction model to provide *T*_*opt*_ estimates for a majority of First, experimentally determined OGTs(*22*) and the 1,438 OGTs predicted with the final OGT SVR model (Figure 2b) were combined to generate a dataset containing the OGT of 22,936 microorganisms. Using this combined dataset 6,507,076 out of 12,115,011 enzymes (54%) in BRENDA could be annotated with the OGT value of their source organism, of which 909,954 enzymes (14%) were contributed by the predicted OGT values. In a second step our *T*_*opt*_ random forest model (Figure 3c) was applied to the 6.5 million OGT annotations combined with the amino acid frequencies of individual enzymes to estimate the *T*_*opt*_. The resulting predictions dramatically added to the *T*_*opt*_ values in BRENDA, increasing them 197-fold (Figure 4a) and covering 4,447 different EC numbers (Figure 4b). Moreover, the temperature coverage, i.e. the minimal and maximal *T*_*opt*_ for an enzyme class, of the vast majority these EC numbers (3,721 of 4,447) were expanded (Figure 4b). The predicted enzyme *T*_*opt*_ and annotated OGT values of these enzymes are freely available for download and re-use (https://zenodo.org/record/2539114, https://doi.org/10.5281/zenodo.2539114).

**Figure 4:**
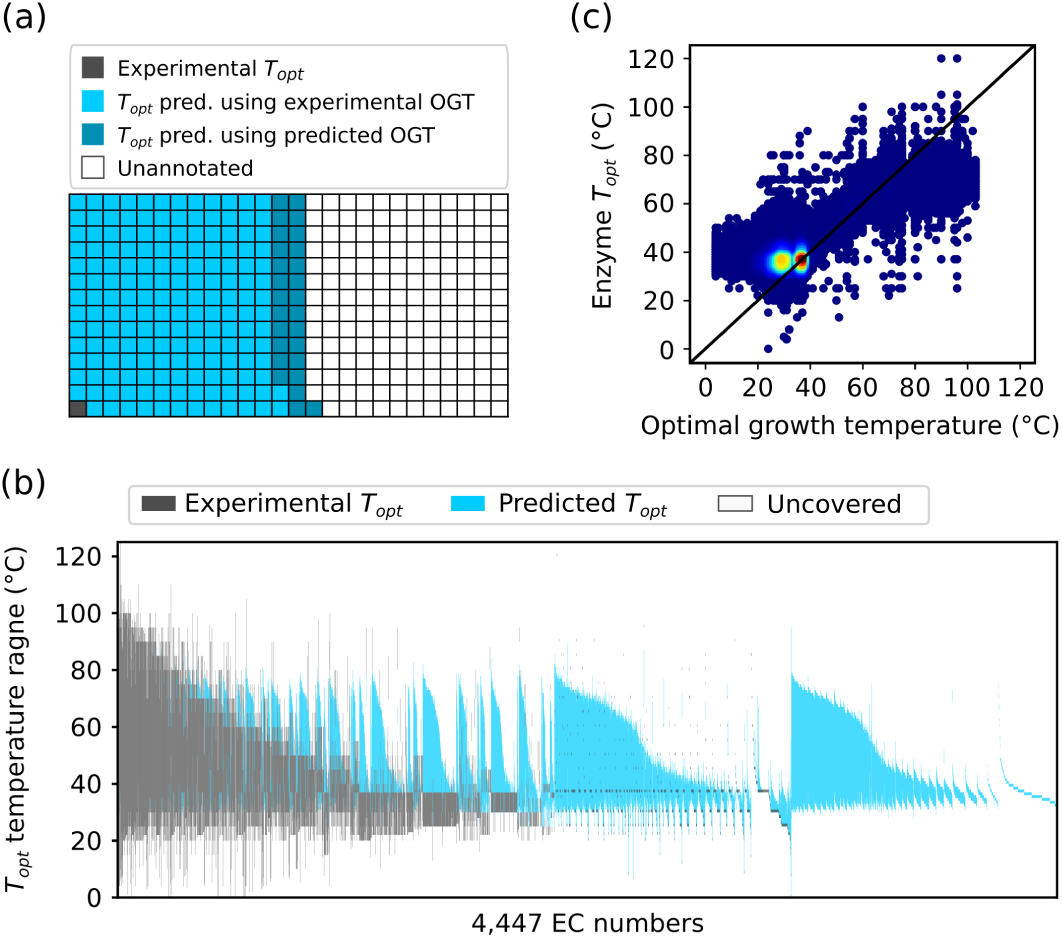
Prediction of enzyme temperature optima. (a) Visual representation of the number of the enzymes with experimental *T*_*opt*_ in BRENDA and the number of enzymes for which *T*_*opt*_ was predicted leveraging experimental and predicted OGTs. Each box represents 33,050 enzymes. There are 12,115,011 enzymes in total. Pred. is an abbreviation of predicted. (b) A visual representation of the *T*_*opt*_ temperature coverage for each EC number after annotation. The span between the highest and lowest *T*_*opt*_ for each enzyme is indicated. Experimental (BRENDA) and predicted *T*_*opt*_ values are shown in different colors. (c) Comparison between OGT of source organism and predicted and experimental *T*_*opt*_ values of enzymes. Colors indicate the density of the points.

As can be seen in Figure 4c, many of the predicted enzyme *T*_*opt*_ values differ significantly from the OGT of the source organism. For enzymes from organisms with OGT below 40°C many have *T*_*opt*_ higher than the OGT. In contrast, enzymes from thermophiles generally have a lower *T*_*opt*_ than the OGT. These results are in good agreement with previous findings comparing experimental OGT of organisms with average enzyme *T*_*opt*_(*22*). For three representative organisms we show that the distribution of predicted *T*_*opt*_ values are indeed consistent with experimental values (Figure S4). The predicted *T*_*opt*_ values provided here represent a rich resource for identifying enzymes suitable for bioprocess carried out at high temperatures.

### 3.6 Tome: a command line tool for OGT prediction and identification of enzyme homologues with different *T*_*opt*_

To ensure easy access to the OGT predictive model for the scientific community, as well as the enzyme data annotated with OGT of their source organism and estimated *T*_*opt*_, we developed the command line tool Tome (<u>T</u>emperature <u>o</u>ptima for <u>m</u>icroorganisms and <u>e</u>nzymes). This tool is simple to use and has two fundamental applications: (1) prediction of OGT from a file containing protein sequences encoded by an organism’s genome; (2) identification of functional homologues within a specified temperature range for an enzyme of interest. For the prediction of OGT, a list of proteomes in fasta format(*50*) is provided as input and the temperature predictions are returned as an output. While this tool will perform predictions on any input given, we stress that the tool has been trained on bacteria, archaea and a only small set of eukarya - mostly fungi and protists. Predictions on organisms which do not fall into these categories may result in inaccurate results. For the identification of enzyme functional homologs with different estimated *T*_*opt*_, one can either simply specify an EC number and temperature range of interest to get all enzyme sequences from BRENDA matching the criteria. Alternatively, the sequence of an enzyme of interest can be provided in fasta format. The algorithm will then perform a protein BLAST(*35*) and an additional output file will be generated containing only homologous enzymes (default e-value cutoff is 10^−10^) within the specified temperature range. Full instructions regarding installation and usage of the Tome tool is available online (https://github.com/EngqvistLab/Tome).

## 4 Author contributions

GL, JN and MKME conceptualized the research. JN acquired funding to support the project. MKME generated the proteome and BRENDA datasets and performed data curation. GL performed the computational and statistical data analysis. GL, KSR and MKME interpreted results. GL wrote the computer code for the Tome package. GL and MKME created the publication figures. GL and MKME wrote the initial draft of the paper. GL, KSR, JN and MKME carried out revisions on the initial draft and wrote the final version.

## Acknowledgement

The computations were performed on resources at Chalmers Centre for Computational Science and Engineering (C3SE) provided by the Swedish National Infrastructure for Computing (SNIC). We thank Pia Schwitters for her assistance with performing the manual literature search for organism growth temperatures.

## Supporting Information Available

Supplemental Figures S1-S4 and Tables S1-S2 are available free of charge on the ACS Publications website at DOI: xx.

## Graphical TOC Entry

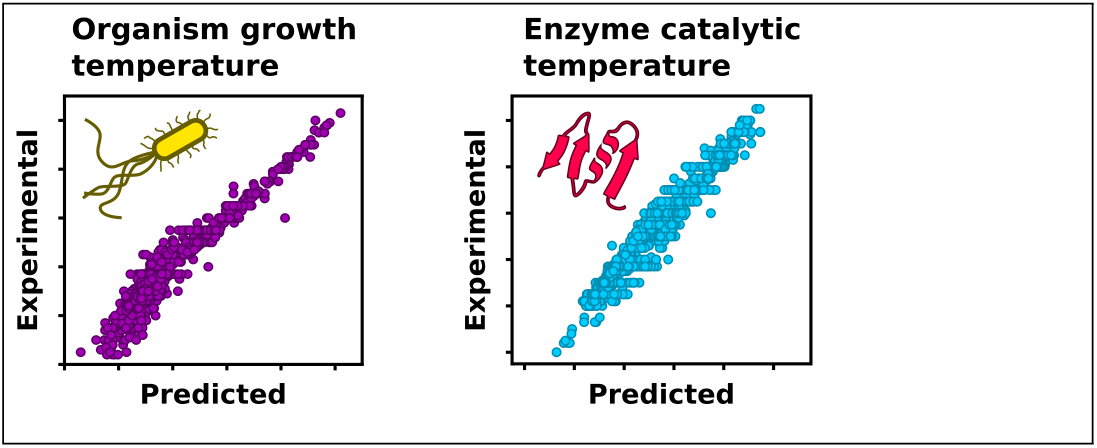

