## Supplementary information for "Machine learning applied to predicting microorganism growth temperatures and enzyme catalytic optima" (from another version of this preprint: version 2, 2019-01-17)

### Supplementary figures

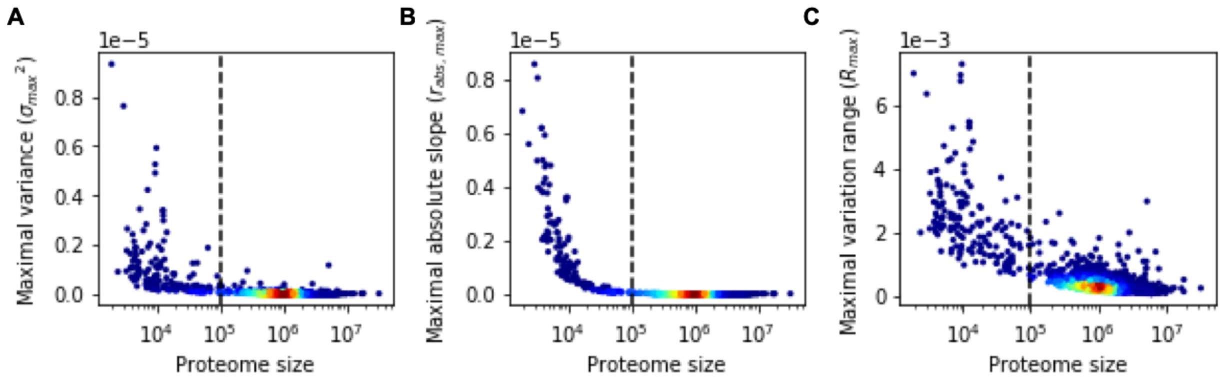


**Supplementary Figure 1 |** The calculated (A) $\boldsymbol{\sigma}^2_{\mathbf{max}}$, (B)$\boldsymbol{r}_{\mathbf{abs,max}}$ and (C) $\boldsymbol{R}_{\mathbf{max}}$ of dipeptides of all species in the dataset were plotted against the number of amino acids in the proteome. These three metrics are the same as defined in Fig. 1b. The dash line indicate the cutoff for the selection of proteomes based on size.

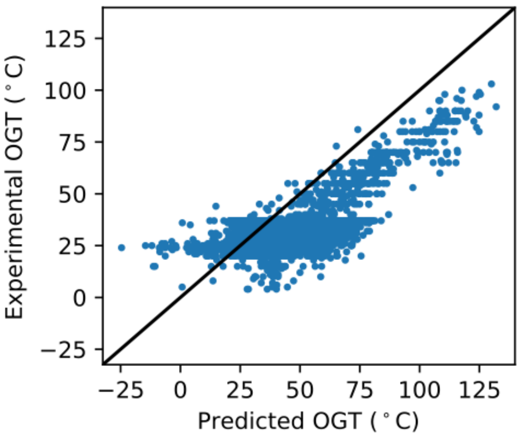


**Supplementary Figure 2 |** The performance of the linear model based on the frequency of seven amino acids I, V, Y, W, R, E, L[12]. The $R^2$ score of the original model is -2.10. If we train the same linear model again with our dataset, an $R^2$ score of 0.48 is achieved.

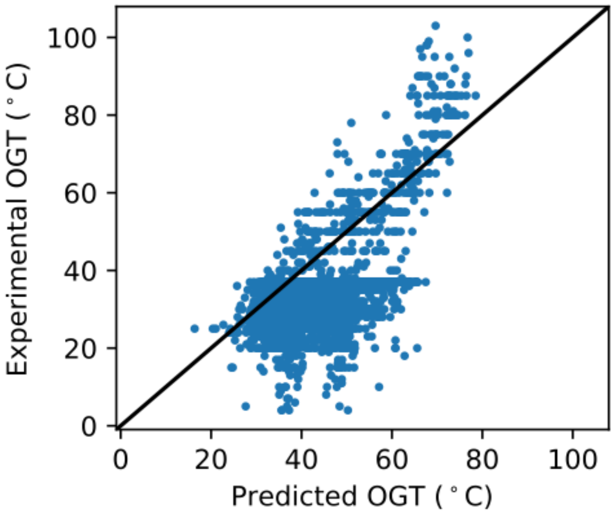


**Supplementary Figure 3 |** The performance of the linear model based on amino acid counts from Nakashima et al.[10] The $R^2$ score of the original model is -0.49. If we train the same linear model again with our dataset, an $R^2$ score of 0.67 is achieved.

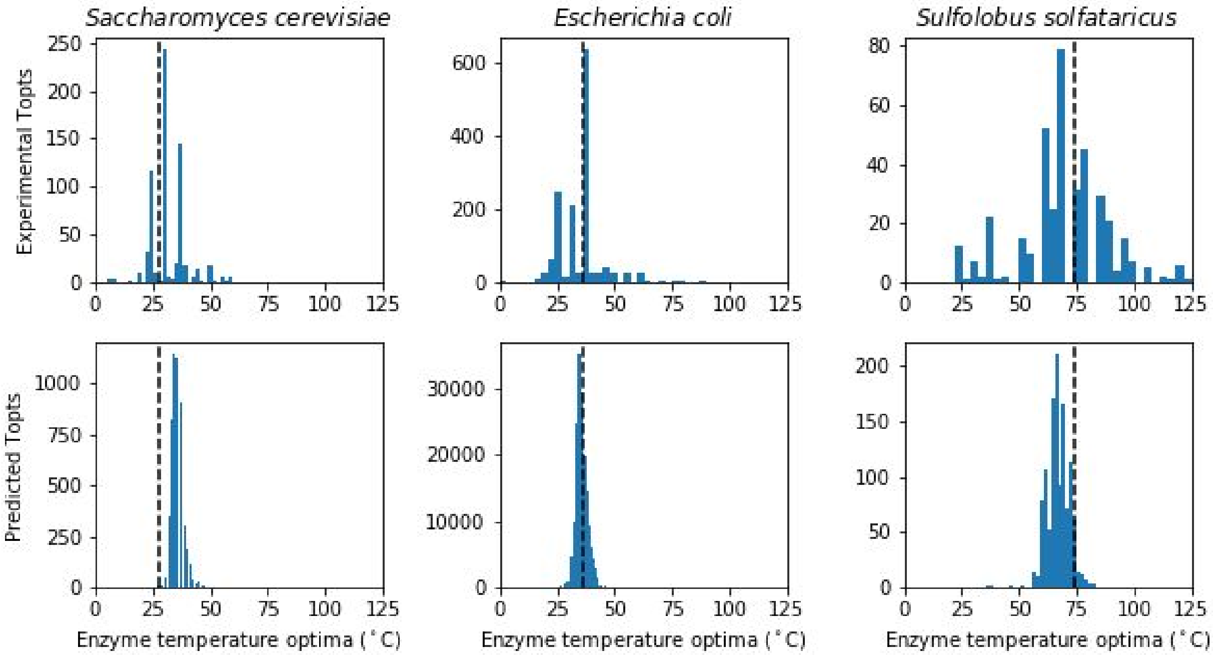


**Supplementary Figure 4 |** Comparison of enzyme $T_{opt}$ distribution between predicted values and experimental values of three microorganisms with rich experimental $T_{opt}$ data in BRENDA. The three histograms in the first row show the distribution of experimental values. The three histograms in the second row show the distribution of predicted values. Each column represent the data from one organism. The dash lines indicate optimal growth temperatures.

### Supplementary Tables

**Supplementary Table 1 | Experimental OGT from literatures for 54 selected organisms**

| Organism | Domain | Growth temperature (°C) | Source |
|---|---|---|---|
| *Oscillochloris trichoides* | Bacteria | 28 | [35] |
| *Gilvimarinus agarilyticus* | Bacteria | 27.5 | [36] |
| *Francisella persica* | Bacteria | 34 | DSMZ (http://www.dsmz.de) |
| *Parageobacillus caldoxylosilyticus* | Bacteria | 55 | [37] |
| *Chryseobacterium tenax* | Bacteria | 20 | [38] |
| *Candidatus midichloria* | Bacteria | NA | |
| *Trichodesmium erythraeum* | Bacteria | 24 | [39] |
| *Thermofilum adornatus* | Archaea | 92 | [40] |
| *Candidatus desulfofervidus* | Bacteria | 60 | [41] |
| *Rhodoluna lacicola* | Bacteria | 20 | DSMZ (http://www.dsmz.de) |
| *Tilletia walkeri* | eukarya | 21 | [42] |
| *Thermofilum uzonense* | Archaea | 85 | DSMZ (http://www.dsmz.de) |
| *Tepidimonas fonticaldi* | Bacteria | 55 | [43] |
| *Thermococcus eurythermalis* | Archaea | 85 | [44] |
| *Thermococcus onnurineus* | Archaea | 80 | [45] |
| *Thermococcus nautili* | Archaea | 87.5 | [46] |
| *Vulcanisaeta thermophila* | Archaea | 85 | [47] |
| *Microcystis panniformis* | Bacteria | 25 | [48] |
| *Thermus amyloliquefaciens* | Bacteria | 65 | DSMZ (http://www.dsmz.de) |

| | | | |
|---|---|---|---|
| *Halapricum salinum* | Archaea | 37 | [49] |
| *Thermococcus piezophilus* | Archaea | 75 | [50] |
| *Pyrococcus yayanosii* | Archaea | 95 | [51] |
| *Agrococcus pavilionensis* | Bacteria | 6 | [52] |
| *Candidatus evansia* | Bacteria | NA | |
| *Desulfurococcus amylolyticus* | Archaea | 90 | DSMZ (http://www.dsmz.de) |
| *Kosmotoga pacifica* | Bacteria | 70 | DSMZ (http://www.dsmz.de) |
| *Aquifex aeolicus* | Bacteria | 95 | [53] |
| *Fervidobacterium thailandensis* | Bacteria | 80 | [54] |
| *Metallosphaera yellowstonensis* | Archaea | 70 | [55] |
| *Acidibacillus ferrooxidans* | Bacteria | 30 | [56] |
| *Vulcanisaeta moutnovskia* | Archaea | 79 | [57] |
| *Rickettsia aeschlimannii* | Bacteria | 32 | [58] |
| *Candidatus methanoperedens* | Archaea | NA | |
| *Mannheimia massilioguelmaensis* | Bacteria | 37 | [59] |
| *Candidatus desulforudis* | Bacteria | 60 | [60] |
| *Halothermothrix orenii* | Bacteria | 60 | [61] |
| *Thermofilum carboxyditrophus* | Archaea | 90 | [62] |
| *Alicyclobacillus mali* | Bacteria | NA | |
| *Sulfolobus islandicus* | Archaea | 77.5 | [63] |
| *Luteipulveratus halotolerans* | Bacteria | 24 | [64] |
| *Thermincola potens* | Bacteria | NA | |

| | | | |
|---|---|---|---|
| *Acetomicrobium hydrogeniformans* | Bacteria | 55 | DSMZ (http://www.dsmz.de) |
| *Kwoniella dejecticola* | eukarya | 27.5 | [65] |
| *Candidatus portiera* | Bacteria | NA | |
| *Streptococcus halotolerans* | Bacteria | 37 | DSMZ (http://www.dsmz.de) |
| *Thermogladius cellulolyticus* | Archaea | 84 | [66] |
| *Stanieria cyanosphaera* | Bacteria | NA | |
| *Thermanaerothrix daxensis* | Bacteria | 60 | DSMZ (http://www.dsmz.de) |
| *Acidianus hospitalis* | Archaea | 80 | [67] |
| *Rhodopirellula sallentina* | Bacteria | 28 | DSMZ (http://www.dsmz.de) |
| *Batrachochytrium dendrobatidis* | eukarya | 25 | [68] |
| *Peptostreptococcaceae bacterium* | Bacteria | NA | |
| *Candidatus kryptobacter* | Bacteria | NA | |
| *Thermus parvatiensis* | Bacteria | 75 | DSMZ (http://www.dsmz.de) |

**Supplementary Table 2 | Regression models**

| Regression model | Module | Hyperparameter range |
|---|---|---|
| Linear model | sklearn.linear_model.LinearRegression | None |
| Elastic net | sklearn.linear_model.ElasticNetCV | Default |
| Bayes ridge | sklearn.linear_model.BayesianRidge | None |
| Support vector regressor | sklearn.svm.SVR | 'C': numpy.logspace(-5, 10, num=16, base=2.0), 'Epsilon': [0, 0.01, 0.1, 0.5, 1.0, 2.0, 4.0] |
| Decision tree | sklearn.tree.DecisionTreeRegressor | 'Min_samples_leaf': numpy.linspace(0.01, 0.5, 10) |
| Random forest | sklearn.ensemble.RandomForestRegressor | 'Max_features': numpy.arange(0.1, 1.1, 0.1) |
